# Dynamics of single-cell protein covariation during epithelial–mesenchymal transition

**DOI:** 10.1101/2023.12.21.572913

**Authors:** Saad Khan, Rachel Conover, Anand R. Asthagiri, Nikolai Slavov

**Author notes:** Data, code & protocols: scp.slavovlab.net/Khan_et_al_2023.

## Abstract

Physiological processes, such as epithelial–mesenchymal transition (EMT), are mediated by changes in protein interactions. These changes may be better reflected in protein covariation within cellular cluster than in the temporal dynamics of cluster-average protein abundance. To explore this possibility, we quantified proteins in single human cells undergoing EMT. Covariation analysis of the data revealed that functionally coherent protein clusters dynamically changed their protein-protein correlations without concomitant changes in cluster-average protein abundance. These dynamics of protein-protein correlations were monotonic in time and delineated protein modules functioning in actin cytoskeleton organization, energy metabolism and protein transport. These protein modules are defined by protein covariation within the same time point and cluster and thus reflect biological regulation masked by the cluster-average protein dynamics. Thus, protein correlation dynamics across single cells offer a window into protein regulation during physiological transitions.

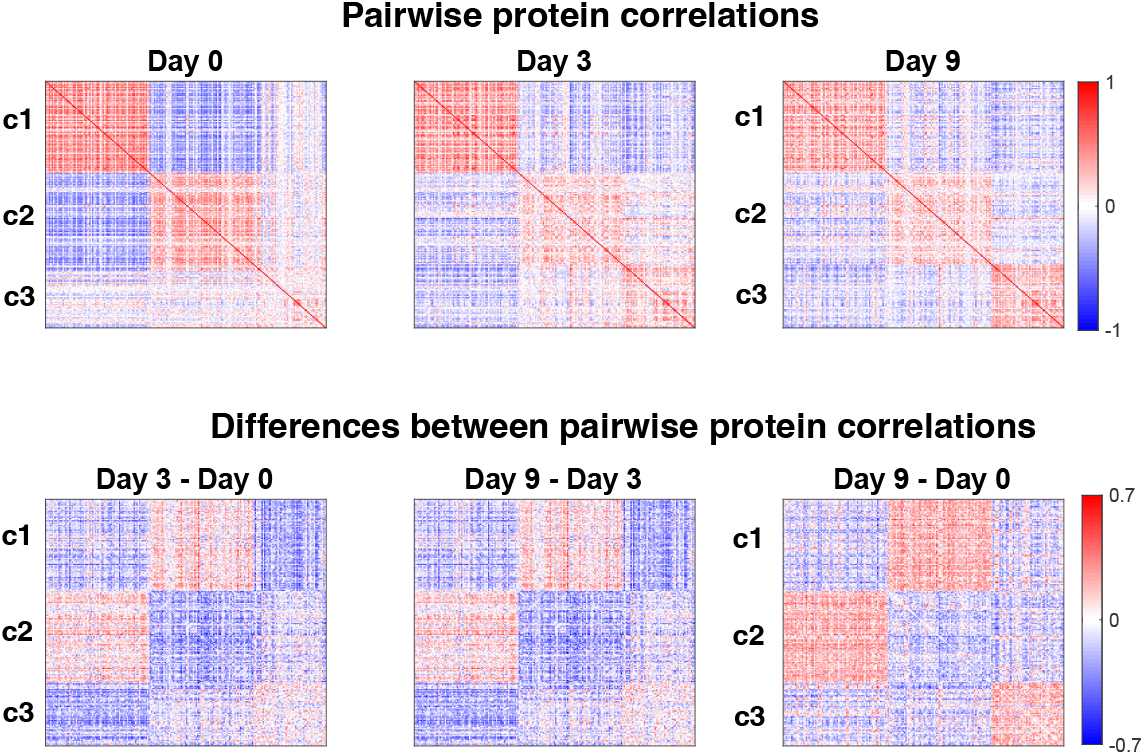

## Introduction

Epithelial-mesenchymal transition (EMT) plays a key role in embryonic and adult development, tissue repair and wound healing, and pathologies such as fibrosis and cancer^1–3^. EMT involves major changes in cell behavior. While most attention is given to cell morphology, adhesion, migration and invasiveness, EMT also impacts cell cycle activity, senescence, apoptosis, metabolism, genomic stability, stemness and drug resistance, among other cell behaviors. The pleiotropic effect of EMT on cell behaviors is mediated by complex regulatory networks, including transcription factors, post-transcriptional and post-translational signaling, intercellular communication and the microenvironment.

EMT regulation across single cells is nonuniform. Single-cell RNA and morphological analysis show significant variability among cells undergoing EMT^4,5^. In what ways variability in protein abundance contributes to single-cell heterogeneity during EMT remains unclear. Since RNA level is an unreliable predictor of protein abundance, protein level variation cannot be reliably inferred from single-cell RNA data^6–9^. Indeed, post-transcriptional processes, such as endocytosis, protein synthesis, modifications and degradation, play a significant role in EMT regulation^10–12^.

While this variability poses challenges for population average measurements, it offers the potential to infer regulatory processes from protein covariation across single cells^13^. Indeed protein co-variation across individual cells may reflect protein interactions and can be detected by singlecell protein measurements with sufficient accuracy, precision and throughput^14,15^. Such analysis is becoming feasible due to the development of powerful single-cell mass spectrometry (MS) proteomic methods^16–24^. As a result, protein covariation across single cells can be quantified^25–28^.

Here, we applied Single Cell ProtEomics^29–31^ to quantify the proteomes of single cells undergoing EMT triggered by TGF*β*, a prominent stimulus for EMT in physiological and pathophysiological processes. Analyzing the system across time, we observed within cluster variation across single cells and strong protein covariation across the single cells within each time point. We systematically quantified this covariation and its change in time. This revealed clusters of proteins whose covariation evolved concertedly in time. These concerted changes in protein covariation include singling proteins, cytoskeleton proteins and metabolic enzymes whose mean abundance per time point does not change. Thus, protein covariation across single cells provides information about cellular remodeling that is complementary to changes in protein abundance.

## Results

### Dynamics of proteins abundance and variability during EMT

We aimed to quantify EMT-mediated changes in protein abundance over the time span of several days. To this end, we performed single-cell proteomics on non-transformed human mammary epithelial cells (MCF-10A) treated with transforming growth factor beta (TGF*β*) for 0 (untreated), 3 and 9 days, Fig. 1a. The chosen dosage of TGF*β* has been shown by us and others to induce overt EMT in MCF-10A cells^32–34^. The time points were chosen to interrogate transient (3 days) and sustained (9 days) responses to TGF*β* during which cells will be in intermediary and overt phases of EMT. Our MS measurements resulted in quantifying about 952 proteins per single cell and 4,571 proteins quantified in at least one cell from the total set of 420 individual cells analyzed, Table S1. However, many of the proteins were quantified in only a fraction of the cells, and we focused on a subset of 1,893 proteins that are quantified in over 30 single cells from the dataset and over 5 single cells from each time point. Such levels of missing data are common in many datasets and limit the number of proteins that can be analyzed without imputing missing values^35^.

**Figure 1.**
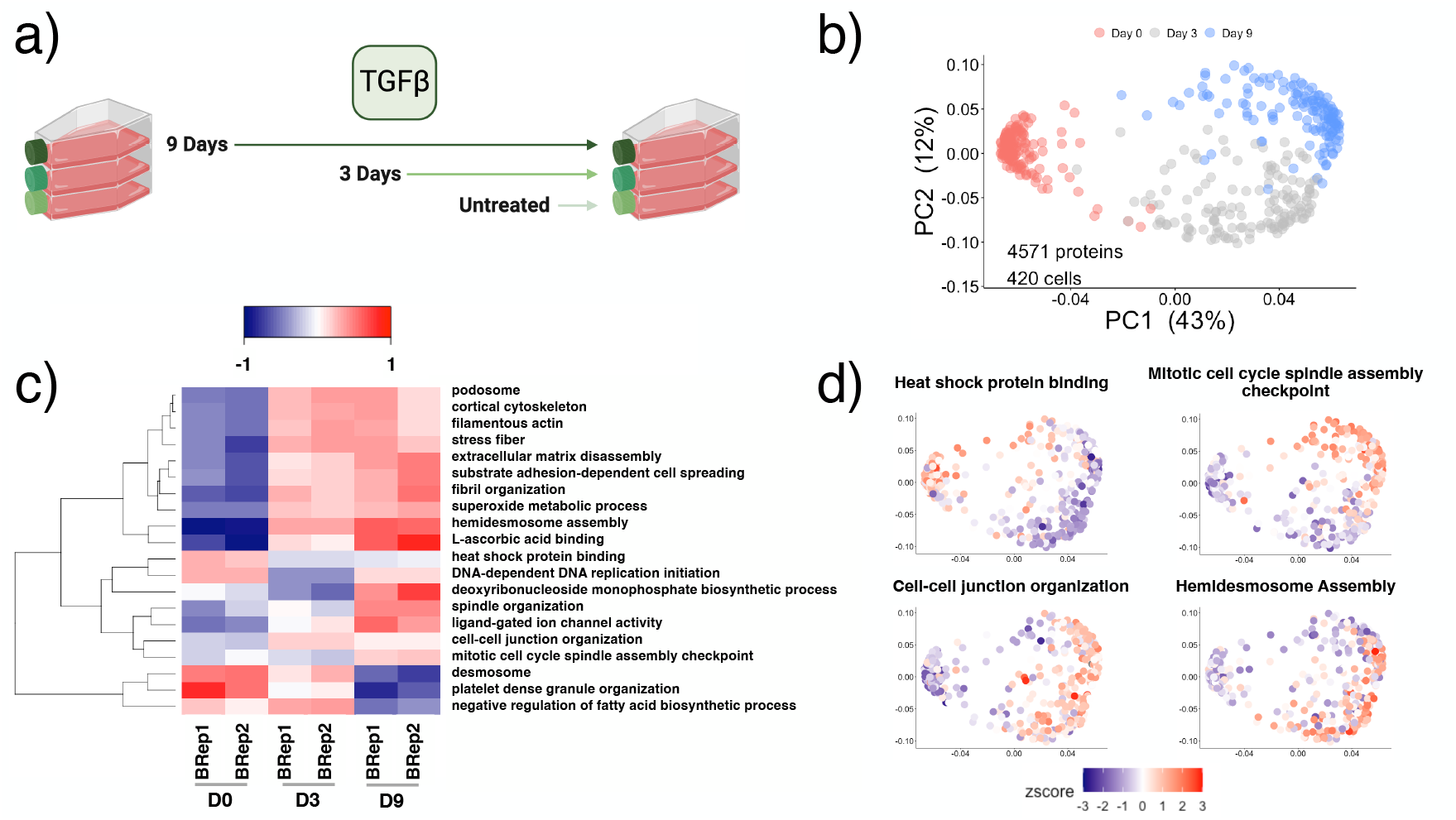
Experimental design, datasets overview and validation. (**a**) Epithelial cells (MCF-10A) were treated by TGF*β* for the indicated duration and then collected for single-cell and bulk protein analysis by MS. (**b**) Single cells plotted in the space of the principal components of their proteome data. The cells are colored by the day post TGF*β* treatment. (**c**) The median abundance of statistically significant protein sets across bulk samples (two biological replicates) analyzed by label-free DIA. (**d**) Single cells in the space of their principal components were colored by the Z scored abundances of select, significantly differential GO terms identified from the bulk proteomes as shown in panel (c).

Principal component analysis (PCA) of protein abundance data shows that cells from the three time points segregate into clusters in PC space, Fig. 1b. The separation suggests that magnitude of changes in protein abundance in time is large enough to readily distinguish cells at different temporal phases of EMT. The first PC primarily separates untreated cells from TGF*β*-treated cells regardless of how far into EMT they have progressed. Cells with transient (3 d) and sustained (9 d) exposure score similarly along PC1 while the second PC separates cells with transient (3 d) and sustained (9 d) exposure to TGF*β*.

Two biological replicates from each time point were analyzed by bulk label-free DIA, and the data analyzed using protein set enrichment analysis^6^. The results indicated multiple functional groups of proteins with differential abundance across the time points, Fig. 1c. Specifically, DNA replication and heat-shock proteins are have higher abundance in epithelial cells, consistent with their active proliferation. Conversely, proteins functioning in cytoskeleton and cell adhesion increase in abundance in days 3 and 9. This trend is particularly strong for hemidesmosome assembly and cell-cell junction proteins. These functional enrichments are highly reproducible across the two biological replicates and generally consistent with previous EMT observations.

To explore the relationship between our bulk and single-cell measurements, we used functional protein groups exhibiting differential abundance in the bulk data to colorcode single cells, Fig. 1d. The results indicate that the protein abundance enrichment is consistent between the bulk and the single-cell measurements, but the single cells exhibit additional variation within a time point. Specifically, we observe within cluster variation around the mean values captured by the bulk measurements. This protein variation across single cells is the focus of our subsequent analyses.

### Dynamics of protein covariation during EMT

Next, we sought to analyze pairwise protein correlations for a set of proteins selected by correlation vector analysis (similar to previously uses in ref^29,36^) to be correlated at both days 0 and 9 but in different ways. An example of such pair is shown in Fig. 2a: RSPH10B and TUBB3 correlate negatively in epithelial cells (Day 0) and positively in mesenchymal cells (Day 9). This change in correlation is statistically significant (p val *<* 10*^−^*^13^, q val *<* 0.1%). Day 3 (whose data was not used for protein selection) exhibits and intermediate correlation pattern, suggesting concerted changes in correlation over EMT progression.

**Figure 2.**
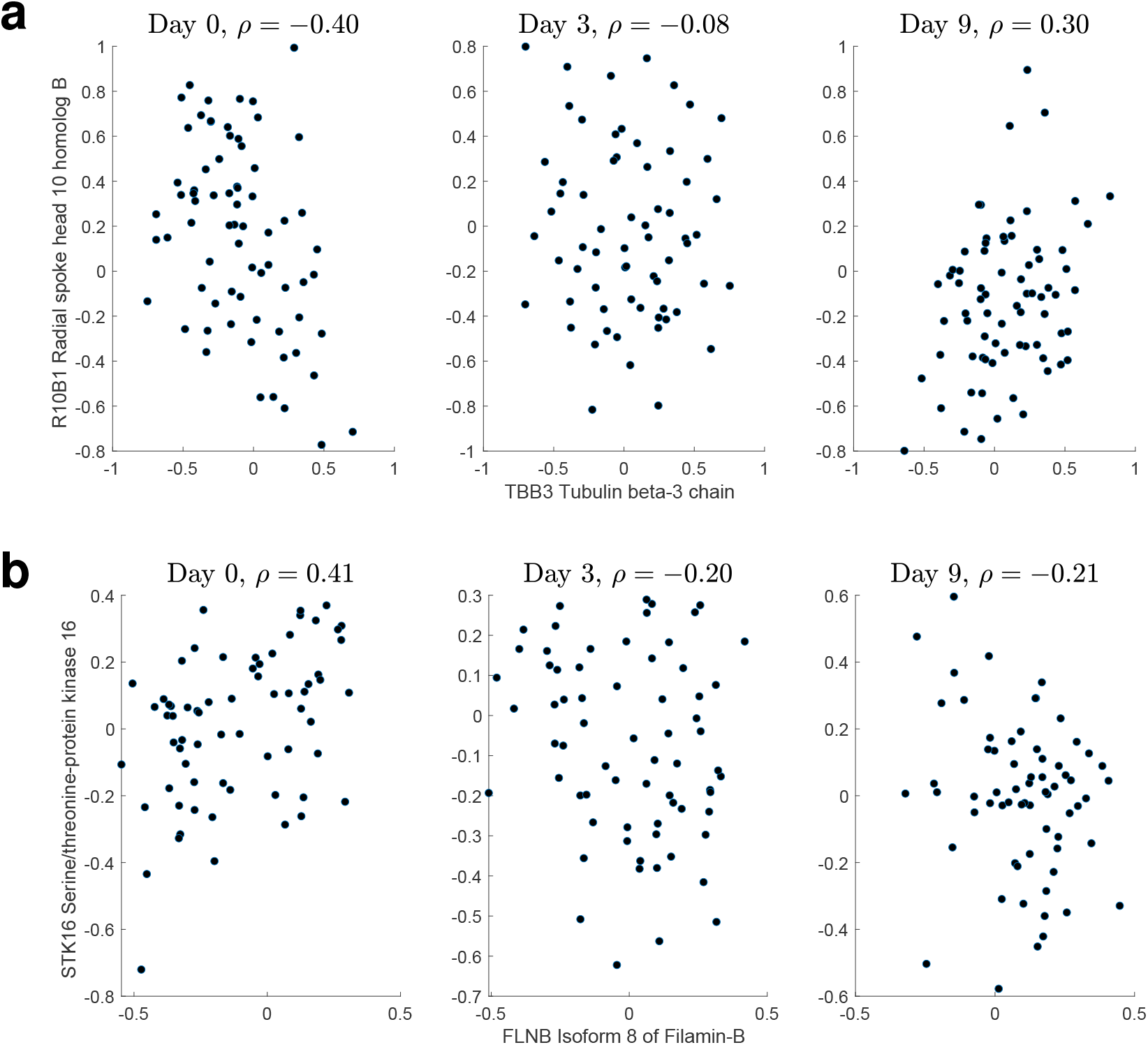
Protein correlations change in the course of EMT. (**a**) Pairwise correlations between RSPH10B and TUBB3 at each of the 3 time points. (**b**) Pairwise correlations between STK16 and FLNB at each of the 3 time points. The probability of observing these changes in correlations in the randomized data is below low, *p <* 10*^−^*^8^, *q <* 10*^−^*^2^.

The change in pairwise correlations in Fig. 2a is not associated with significant changes in the mean abundance of these proteins in Days 0, 3 and 9, suggesting that the changes in correlation patterns may reveal molecular rearrangements inaccessible from measuring mean protein levels.

This observation is bolstered by the data for other proteins pairs exhibiting similarly concerted temporal changes in covariation without significant changes in mean abundance, Fig. 2b.

Next, we sought to expand the correlation patterns suggested by individual pairs of proteins (Fig. 2) to a systematic exploration of changing global patterns of covariation, Fig. 3. To this end, we started by selecting the subset of proteins with multiple pairwise observations (i.e., all proteins for which we can compute pairwise correlations from measured protein abundances) and substantial changes in covariation. To identify proteins with changing correlations, we subtracted the correlations for Day 0 from the correlations for Day 9 and computed the norms of the vectors of correlation differences. Then we selected the proteins (50% percentile) having the largest magnitude difference between Day 0 and Day 9 (Fig. S1) or (20% percentile) having the largest magnitude difference between Day 0 and Day 9 (Fig. S2). The correlations between these proteins defined 3 main clusters when clustered hierarchically Fig. S1, and thus we used K-means clustering with k=3 to define 3 discrete clusters (c1, c2, and c3).

**Figure 3.**
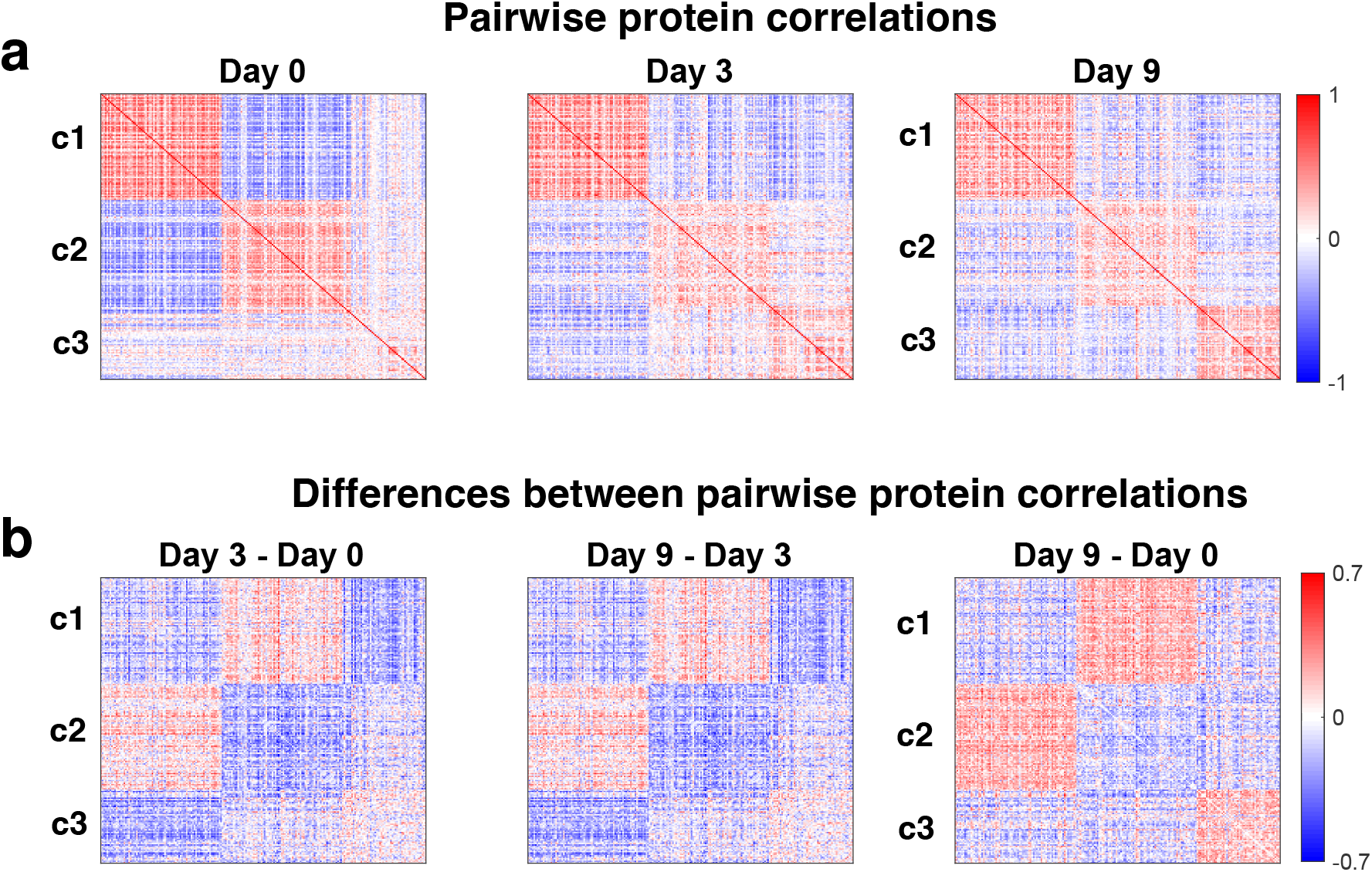
Dynamics of protein covariation during EMT. (**a**) Matrices of pairwise protein correlations at days 0, 3 and 9 clustered using k-means clustering with *k* = 3. The 3 clusters are denoted by c1, c2, and c3. (**b**) Matrices of differences between pairwise protein correlations for the indicated time points. The rows and columns for all days correspond to the same proteins ordered in the same way, namely based on clustering the matrix of correlation differences between Day 9 and 0.

These 3 clusters are well defined, as shown by the matrices of pairwise correlations within each cluster (Fig. 3a) and their corresponding differences across time (Fig. 3b). The global change in correlations for each cluster is monotonic (Fig. 3), and this monotonicity is quantified by the dynamics of mean cluster correlations displayed in Fig. 4a. Since the data from Day 3 have not been used for selecting or clustering proteins, their consistency with the monotonic trends bolsters our confidence in the results.

**Figure 4.**
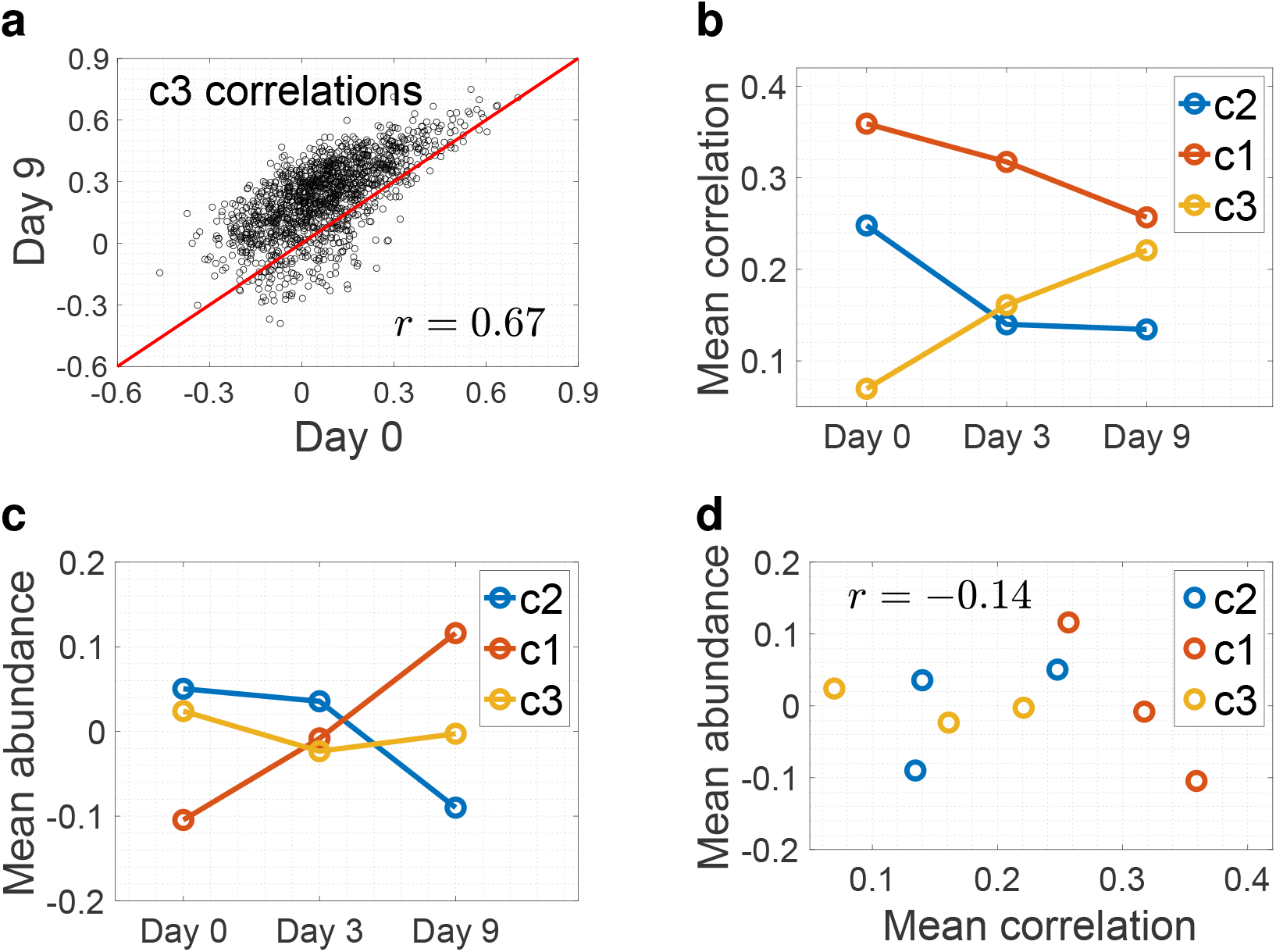
Differences between the dynamics of within-cluster protein correlations and cluster-average abundance. (**a**) Comparison of pairwise correlations for cluster 3 (c3) between days 0 and 9. (**b**) Mean correlations among the proteins from each cluster across time. (**c**) Mean abundance of the proteins from each cluster across time. (**d**) Correspondence between mean cluster correlations and mean cluster protein abundances. The correlation is weak (*r* = *−*0.14) and not statistically significant (*p* = 0.7).

Remarkably, the dynamics of the mean protein correlations for clusters 1,2, and 3 are not reflected in the corresponding dynamics of mean protein abundances, Fig. 4. For a particular cluster, such as c3, the correlations scale in time while the proteins from the cluster remain positively correlated to each other, as shown in Fig. 4a. Based on this observation, we estimated the mean correlation for each cluster and plotted the estimates over time, Fig. 4b. Similarly we estimates, the mean protein abundance of each cluster over time (Fig. 4c), and evaluated the dependence between these mean cluster estimates as shown in Fig. 4d. The result indicates no dependence between the dynamics of protein correlations and mean protein abundance, consistent with the observation for the protein pairs shown in Fig. 2. There results reveal biological signals reflected in the protein covariation across single cells from the same time point but not from the corresponding cluster-average protein abundance, Fig. 4a-c.

To identify biological functions corresponding to each cluster, we performed gene ontology (GO) term analysis^37^ for terms enriched in each cluster relative to all analyzed proteins. We found a many statistically significant protein groups in these clusters, which are provided as Supplemental Tables and a few characteristic groups are highlighted in Table 1. The first cluster comprises proteins involved in cytsokeletal regulation, including actin, vimentin, tubulin subunits, vinculin, filamins and contractility regulators, such as RhoA, myosin and tropomyosin. The proteins in this cluster broadly span cytoskeletal regulation and show statistically significant enrichment when the analysis is extended to more proteins, Fig. S2. Meanwhile, the proteins in the second and third clusters showed statistically-significant enrichment for metabolism and protein synthesis and transport, respectively (Supplemental Tables). The second cluster was enriched for proteins involved in glycolysis and oxidative phosphorylation, consistent with the role of EMT in regulating aerobic and anaerobic utilization of glucose^38^. Many of the enriched functions found in the third cluster involved protein synthesis and transport, including core ribosomal proteins. The third cluster is enriched for other functions associated with EMT, including telomere regulation, and senescence and cell response to DNA damage. Taken together, the protein functions found across the three clusters correspond to the multi-faceted effect of EMT on cellular functions.

**Table 1.** Enriched biological functions in the correlation clusters. The table summarizes protein sets and associated representative proteins form performing GO Gorilla analysis on cluster c1, c2, and c3. All terms are significant at 1% FDR. The full results from the analysis are available as Supplemental Tables.

| Cluster | Enriched function | Proteins |
| --- | --- | --- |
| c1 | regulation of actin filament-based process | fascin, RhoA, IQGAP1, Arp3 |
| c2 | glucose and pyruvate metabolic processes<br>oxidative phosphorylation | pyruvate kinase, aldolase, enolase, lactate dehydrogenase<br>ATP synthase, cytochrome C oxidase |
| c3 | protein synthesis and transport<br>telomere maintenance, cell response to DNA damage | ribosomal proteins<br>XRCC5, XRCC6, PARP1, PCNA |

## Discussion

Protein covariation across single cells may identify regulatory interactions^13^, and here we demonstrate its potential to delineate dynamic remodeling of protein networks during EMT. Our work builds upon previous observations of RNA covariation^39^ and protein covariation^25–28^ and extends the anaysis and interpretation towards temporal dynamics during cellular transitions.

Proteins covariation has to important benefits relative to RNA covariation for inferring biological regulation. First, proteins quantification is based on sampling sufficient number of copies from most proteins (often 100s of copies)^29,40,41^ to support reliable quantification of correlations across single cells (not clusters of cells), as shown in Fig. 2. Second, much of the regulation may be driven by protein synthesis and degradation of protein subunits of complexes^42^, and this component is detectable only at the protein level. For these two reasons, protein covariation offers an informative perspective towards biological regulation^13^.

We based our analysis on correlations between proteins with many pairwise observations. Many of the proteins quantified in our dataset did not have sufficient number of pairwise observations to be included in this analysis due to the stochastic approach of shotgun data acquisition. This limitations can be overcome in future studies by using prioritised data acquisition (pSCoPE)^28,43^ or multiplexed data independent acquisition (plexDIA)^44–46^. Thus, using the latest generation of single-cell proteomic technology and future innovations^47^ will further empower the approach that we used in this study.

Comparison of mean abundance and covariation over the time course of EMT provides intriguing insights. For cluster 1 (cytoskeletal proteins), the mean abundance of proteins increases, as generally expected during EMT progression. However, the correlation in the expression of cytoskeletal regulators decreases, suggesting that the expression of cytoskeletal regulators becomes more heterogeneous in the population as EMT progresses. Since the cytoskeleton confers cell shape, one predicts that increased heterogeneity in its regulators will lead to cell morphological variability. This expectation is consistent with single-cell analysis of cell morphology that shows increased heterogeneity in cell shape as EMT progresses^48^.

In addition to cell shape changes, EMT shifts the balance between aerobic and anaerobic utilization of glucose^38^. However, our data show that the mean abundance of cluster 2 proteins (metabolism) remains constant during EMT progression. Thus, changes in mean abundance do not explain shifts in metabolism. In contrast, the correlation in expression of cluster 2 proteins increases as EMT progresses, suggesting that functional changes in metabolism may be achieved through coordinated changes in abundance (covariation) rather than change in mean abundance.

## Materials and methods

### Cell culture

Non-transformed human mammary epithelial cells (MCF-10A, ATCC) were maintained in growth medium consisting of DME medium/Ham’s F-12 (ThermoFisher) containing HEPES and L-glutamine supplemented with 5% (v/v) horse serum (ThermoFisher), 20 ng/ml EGF (Peprotech), 0.5 *µ*g/ml hydrocortisone, 0.1 *µ*g/ml cholera toxin, 10 *µ*g/ml insulin (Sigma), and 1% penicillin-streptomycin (ThermoFisher), as described previously^49^. To induce EMT, cells were treated with 20 ng/ml TGF*β* (Peprotech) in growth medium for 0, 3 and 9 days^32^. TGF*β*-containing medium was refreshed every three days.

### Sample preparation

Cells were harvested as single-cell suspension and prepared for MS analysis using Nano-ProteOmic sample Preparation (nPOP) as described by Leduc *et al.*^50,51^. The automated collection of prepared samples had not been developed yet^26^ and so samples were manually collected using a pipette (using 5*µl* of mass spectrometry grade Acetonitrile then water respectively) and transferred into a 384-well plate (Thermo AB1384). The samples were then dried down in a SpeedVac vacuum evaporator and resuspended in 1.07*µl* of 0.1% Formic Acid (buffer A) and tightly sealed using an aluminium foil cover (Thermo Fisher AB0626).

For the bulk experiments, cells were harvested (in MS grade water, at roughly 2000 cell/*µl*) and frozen at −80C. The samples were prepared using mPoP^52^, following guidelines for the digest of carriers as outlined in Petelski *et al.*^30^. Post digest, the samples were dried down in a SpeedVac vacuum evaporator and resuspended at a concentration of 1 *µg*/*µl* in 0.1% Formic Acid (buffer A) in a glass insert with polyspring within an HPLC vial.

### Peptide separation and MS data acquisition

The separation of the single cell samples was performed at a constant flow rate of 200nL/min using a Dionex UltiMate 3000 UHPLC. From the 1.07*µl* of sample in each well, 1 *µl* was loaded onto a 25cm *×* 75 *µM* IonOpticks Odyssey Series column (ODY3-25075C18). The separation gradient was 4% buffer B (80% acetonitrile in 0.1% Formic Acid) for 11.5 minutes, a 30 second ramp up to 8%B followed by a 63 minute linear gradient up to 35%B. Subsequently, buffer B was ramped up to 95% over 2 minutes and maintained as such for 3 additional minutes. Finally, buffer B was dropped to 4% in 0.1 minutes and maintained for 19.9 additional minutes.

The mass spectra were acquired using a Thermo Scientific Q-Exactive mass spectrometer from minutes 20 to 95 of the LC method. The electrospray voltage of 1700 V was applied at the liquid liquid junction of the analytical column and transfer line. The temperature of the ion transfer tube was 250°C, and the S-lens RF level was set to 80.

For the single cell samples, after a precursor scan from 450 to 1600 m/z at 70,000 resolving power, the top 7 most intense precursor ions with charges 2 to 4 and above the AGC min threshold of 20,000 were isolated for MS2 analysis via a 0.7 Th isolation window with a 0.3 Th offset. These ions were accumulated for at most 300 ms before being fragmented via HCD at a normalized collision energy of 33 eV (normalized to m/z 500, z = 1). The fragments were analyzed at 70,000 resolving power. Dynamic exclusion was used with a duration of 30 s with a mass tolerance of 10 ppm.

The bulk sample separation and mass spectra acquisition was carried out using the V2 method outlined by Derks *et al.*^44^, briefly, the duty cycle for the data independent acquisition of spectra consisted of an MS1 scan at 70,000 resolving power limited to a maximum injection time of 300ms, an AGC maximum of 3 × 10^6^ and normalized collision energy of 27. Each MS1 scan was followed by 40 MS2 scans at 35,000 resolving power, an AGC max of 3 × 10^6^ and a maximum fill time of 110ms. The DIA window widths, in order, were: 25 × 12.5 Th, 7 × 25Th and 8 × 62.5 Th, there was a 0.5Th overlap in windows.

### MS data searching

The raw single cell data was searched by MaxQuant^53^, a software package for proteomics data analysis, against a protein sequence database that included all entries from the human SwissProt database and known contaminants. The MaxQuant search was performed using the standard workflow, which includes trypsin digestion and allows for up to two missed cleavages for peptides with 7 to 25 amino acids. Tandem mass tags (TMTPro 16plex) were specified as fixed modifications, while methionine oxidation and protein N-terminal acetylation were set as variable modifications. Carbamidomethylation was disabled as a fixed modification because it was not performed. Second peptide identification was also disabled. The calculation of peak properties was enabled. All peptide-spectrum-matches (PSMs) and peptides found by MaxQuant were exported to the evidence.txt files. The confidence in the PSMs was further updated using DART-ID, which is a Bayesian framework for increasing the confidence of PSMs that were consistently identified at the same retention time with high-confidence PSMs for the same amino acid sequences^54^. The updated data were filtered at 1% FDR for both peptides and proteins as described by Petelski *et al.*^30^.

The bulk data was searched using DIANN^55^ version 1.8.1 beta 7 using a spectral library that was prepared via an in-silico digest of a Swissprot Fasta database that contained 20,375 proteins. The mass accuracy was set to 10, methionine excision was set as a variable modification, MBR was on and outputs filtered at 1% FDR.

### Data filtering and analysis

The peptide by cells matrices were processed by the SCoPE2 analysis pipeline^29,30,56^, which resulted in 4,571 proteins quantified across 420 single cells. However, each single cell had only about 1,000 proteins quantified and many proteins were quantified in relatively few single cells. Thus, we subset the proteins to the 1,893 proteins quantified in at least 30 single cells from the dataset and at least 3 single cells from each time point.

We computed the pairwise Pearson correlations for each time point between these 1,893 proteins using only measured abundances, without imputation, which resulted in three 1, 893 *×* 1, 893 correlation matrices, *R*_1_, *R*_2_, and *R*_3_ for days 0, 3 and 9 respectively. We further computed the Pearson correlations among correlation vectors of *R*_1_ and *R*_3_ corresponding to the same protein as previously described^25,29,36^, resulting in a vector **v**. Each element of **v** corresponds to one protein and quantifies the similarity of its correlations to the remaining 1,892 proteins. The elements of **v** were used to explore pairwise combination of proteins whose correlations change significantly between days 0 and 9 and the examples shown in Fig. 2. To calculate the statistical significance for the change in the correlations, we computed the same correlation difference for 10^8^ randomized samples and estimated the fraction of randomized samples whose correlation difference exceeds the difference observed in the data.

For the systematic analysis of correlations in Fig. 3, we computed the matrix of correlation differences Δ*R* = *R*_3_ *− R*_1_ and preserved only its rows and columns corresponding to a set *ϕ* of 418 proteins with no missing data, i.e, the corresponding correlations could be computed only from quantified proteins both for day 0 and 9. Then, we quantified the average magnitude of correlation change for each of these 418 proteins by computing the norm of its corresponding column in Δ*R_ϕ_*, resulting in a vector **m**. To select the proteins whose correlations change the most, we selected the subset *ω* of 209 proteins having norms in **m** larger than the 50% percentile of **m** (the median of **m**). Then we performed means clustering with *k* = 3 on Δ*R_ω_*and used the 3 resulting clusters to display *R*_1_*_ω_*, *R*_2_*_ω_*, and *R*_3_*_ω_* and their differences in Fig. 3. As an alternative approach, we clustered Δ*R_ω_* hierarchically and used the resulting permutation to order the rows and columns of *R*_1_*_ω_*, *R*_2_*_ω_*, and *R*_3_*_ω_* and their differences, as displayed in Fig. S1.

To quantitatively display the dynamics of correlations for the 3 clusters derived by K-means clustering, we computed the mean correlation for all pairwise correlations between proteins assigned to a cluster for each of the 3 time points, Fig. 4. Similarly, we computed the average protein abundance for all proteins (log2 fold change relative to the mean) assigned to a cluster for each of the 3 time points. Only measured protein values were used for computing the average-cluster protein abundance; no imputation values were used.

For the bulk samples, DIA-NN reports were further filtered at 1% Lib.PG.Q.Value and subset to contain only proteotypic peptides. The MS2 level peptide intensities (Precursor.Normalised) were collapsed to protein levels across runs using the diannMaxLFQ function from the ‘diann’ R package. Each bulk samples protein level outputs were normalised to its median to account for differential loading, converted to relative levels using the mean across runs and finally log2 transformed.

Protein set enrichment analysis (PSEA) was subsequently carried out for each biological replicate individually. The Kruskall-Wallis test was used to test the significance of the difference in relative intensity distributions of proteins belonging to a GO term across the three time points. Only GO terms where at least 30% of proteins were present across samples were tested and the maximum number of proteins per GO term was limited to 55. The p values obtained across all tests were adjusted for the multiple hypotheses tested using the Benjamini Hochberg procedure and resultant Q values were used to control the FDR at 5%. The median value of proteins within a GO term was used to represent its relative abundance.

## Supporting information

Supplemental Tables-clusters-GOrilla

## Availability

Data, metadata, code, and protocols are organized according to community recommendations^24^ and available at supplemental information and at scp.slavovlab.net/Khan_et_al_2023. The Raw MS data are available at MassIVE: MSV000092872 and ProteomeXchange: PXD045423.

## Acknowledgments

We thank Morgan Benson, Audrey Kidd and Andrew Leduc for help with sample preparation and members of the Asthagiri and Slavov labs for feedback and helpful discussions. The work was funded by an NCI award R21CA246150-01A1 to A.R.A. and N.S., and Allen Distinguished Investigator award through The Paul G. Allen Frontiers Group to N.S., an NIGMS award R01GM144967 to N.S., and a MIRA award from the NIGMS of the NIH to N.S. (R35GM148218) to N.S.

## Supplemental Figures

**Table S1.** Overview of Protein identifications. The table summarizes the number of proteins identified over different levels of data completeness (filtered at 1% FDR)^29,30^.

| Criteria | In at least 1 cell | Median / cell | In at least 30 cells | Pairwise |
| --- | --- | --- | --- | --- |
| # Proteins | 4,571 | 952 | 1,893 | 418 |

**Figure S1.**
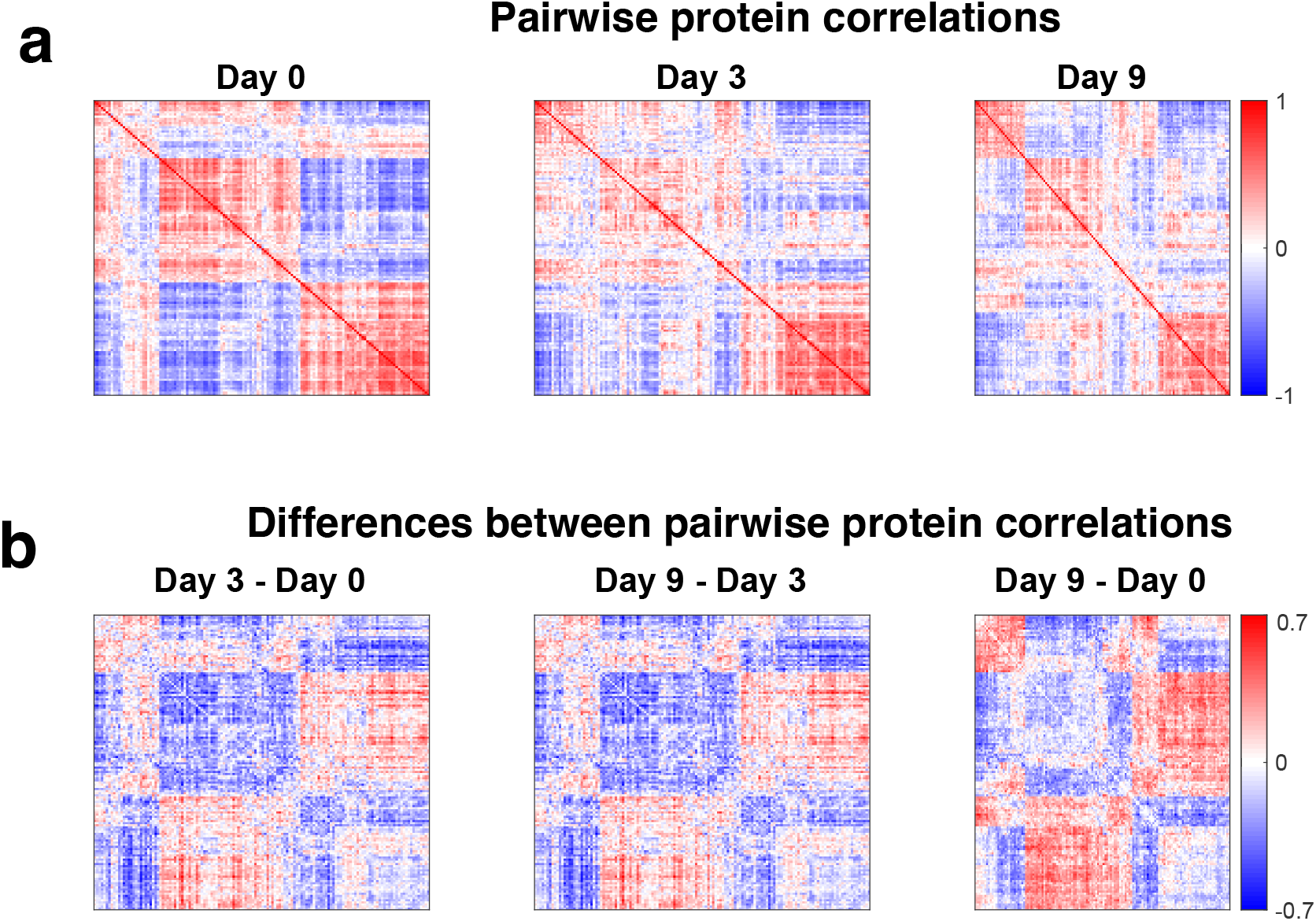
Dynamics of protein covariation during EMT. The the proteins whose correlations have the largest magnitude (exceeding the 50% percentile) difference between Day 0 and Day 9 were selected and their correlation matrices clustered hierarchically. (**a**) Matrices of pairwise protein correlations at days 0, 3 and 9. (**b**) Matrices of differences between pairwise protein correlations for the indicated time points. The rows and columns for all days correspond to the same proteins ordered in the same way, namely based on clustering the matrix of correlation differences between Day 9 and 0.

**Figure S2.**
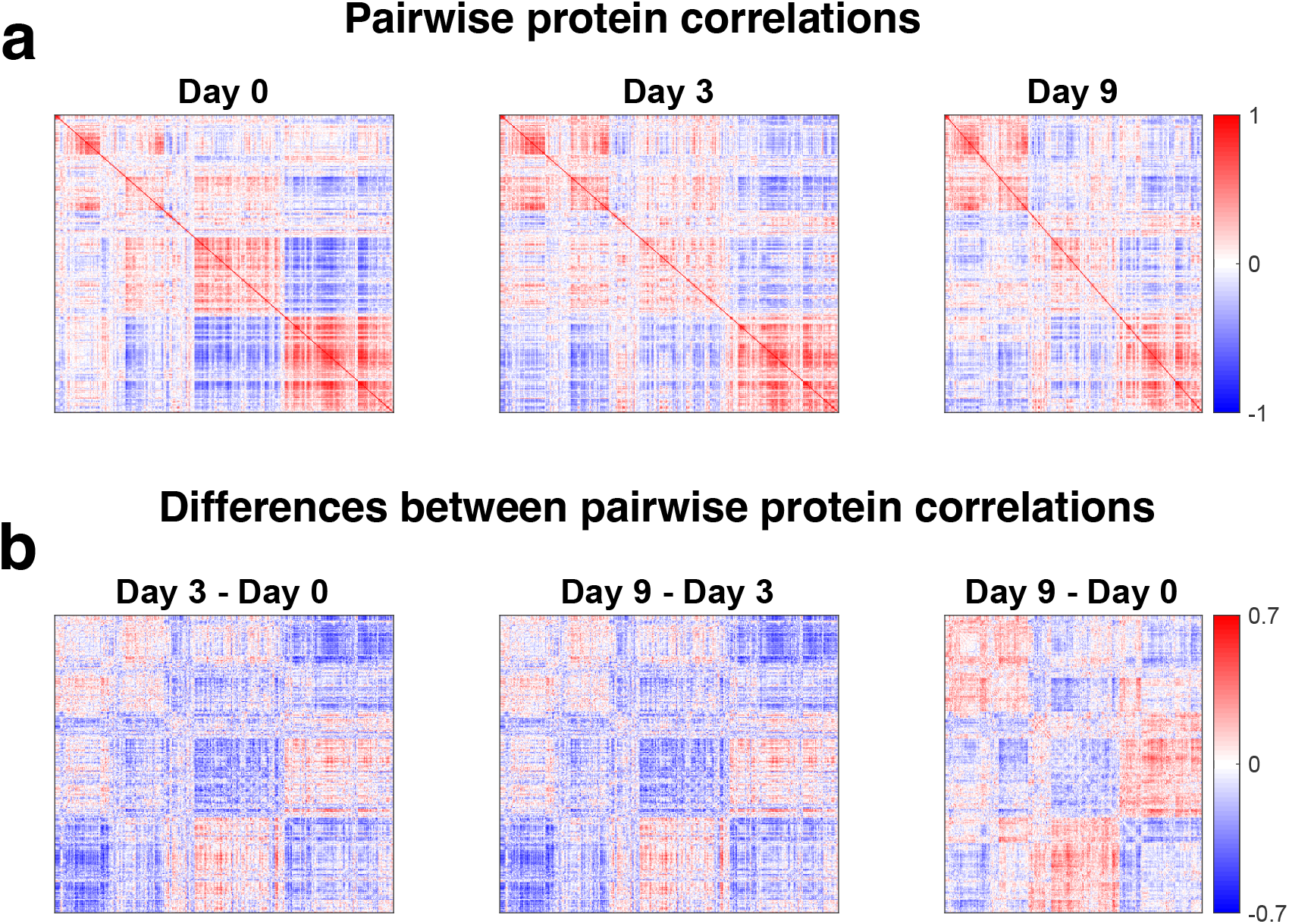
Dynamics of protein covariation during EMT. The the proteins whose correlations have large magnitude (exceeding the 20% percentile) difference between Day 0 and Day 9 were selected and their correlation matrices clustered hierarchically. (**a**) Matrices of pairwise protein correlations at days 0, 3 and 9. (**b**) Matrices of differences between pairwise protein correlations for the indicated time points. The rows and columns for all days correspond to the same proteins ordered in the same way, namely based on clustering the matrix of correlation differences between Day 9 and 0.

