## Supplementary figures and images for "Dynamics of single-cell protein covariation during epithelial–mesenchymal transition"

### Cluster-1.png

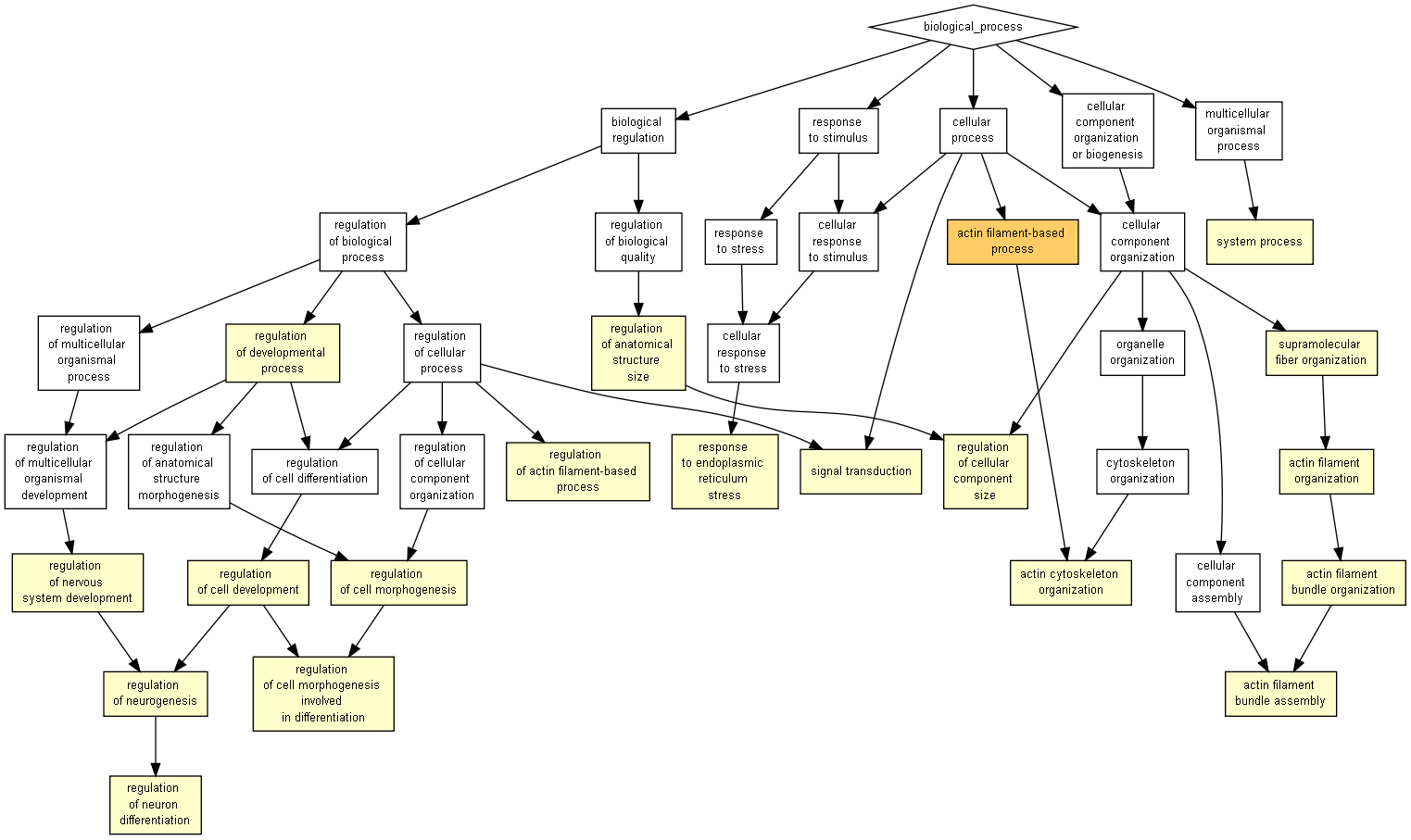

### Cluster-2.png

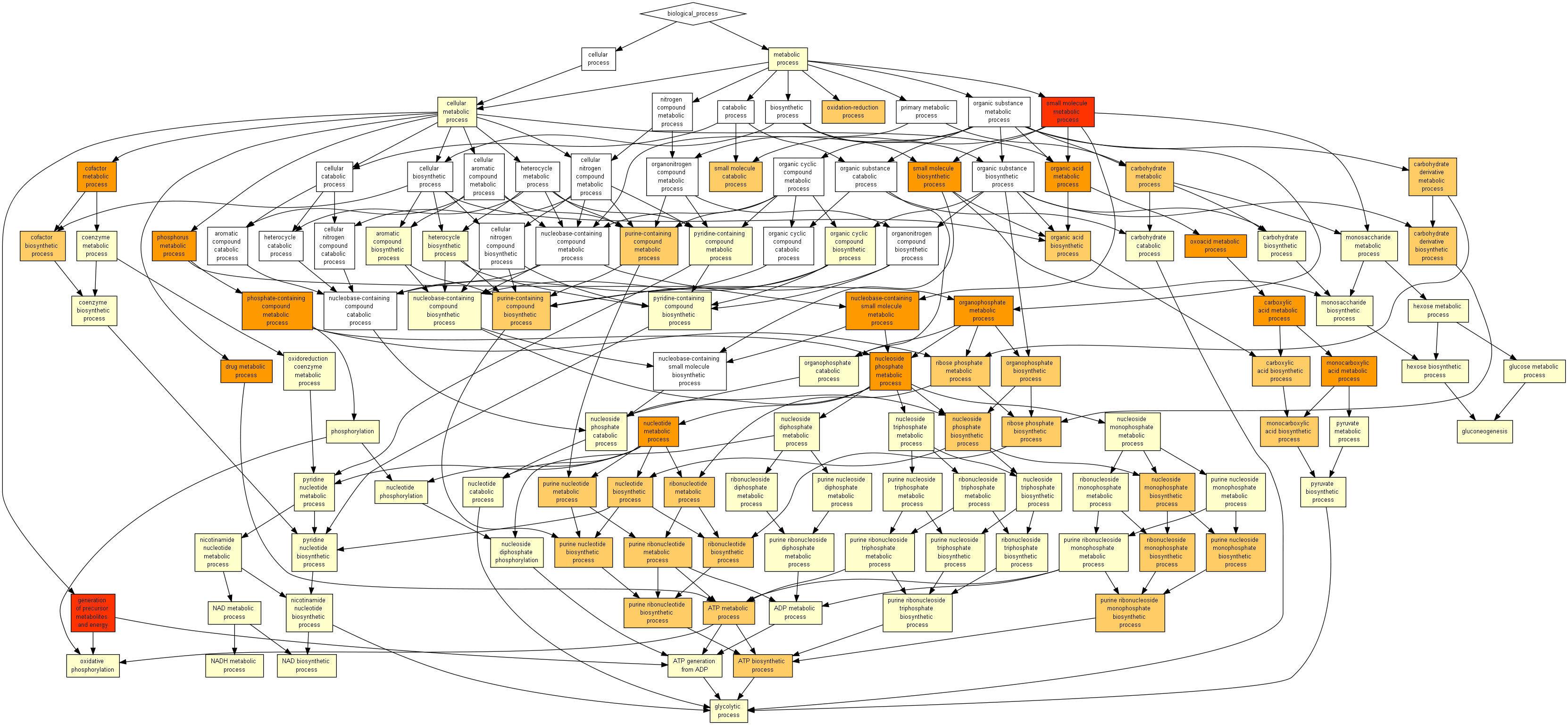

### Cluster-3.png

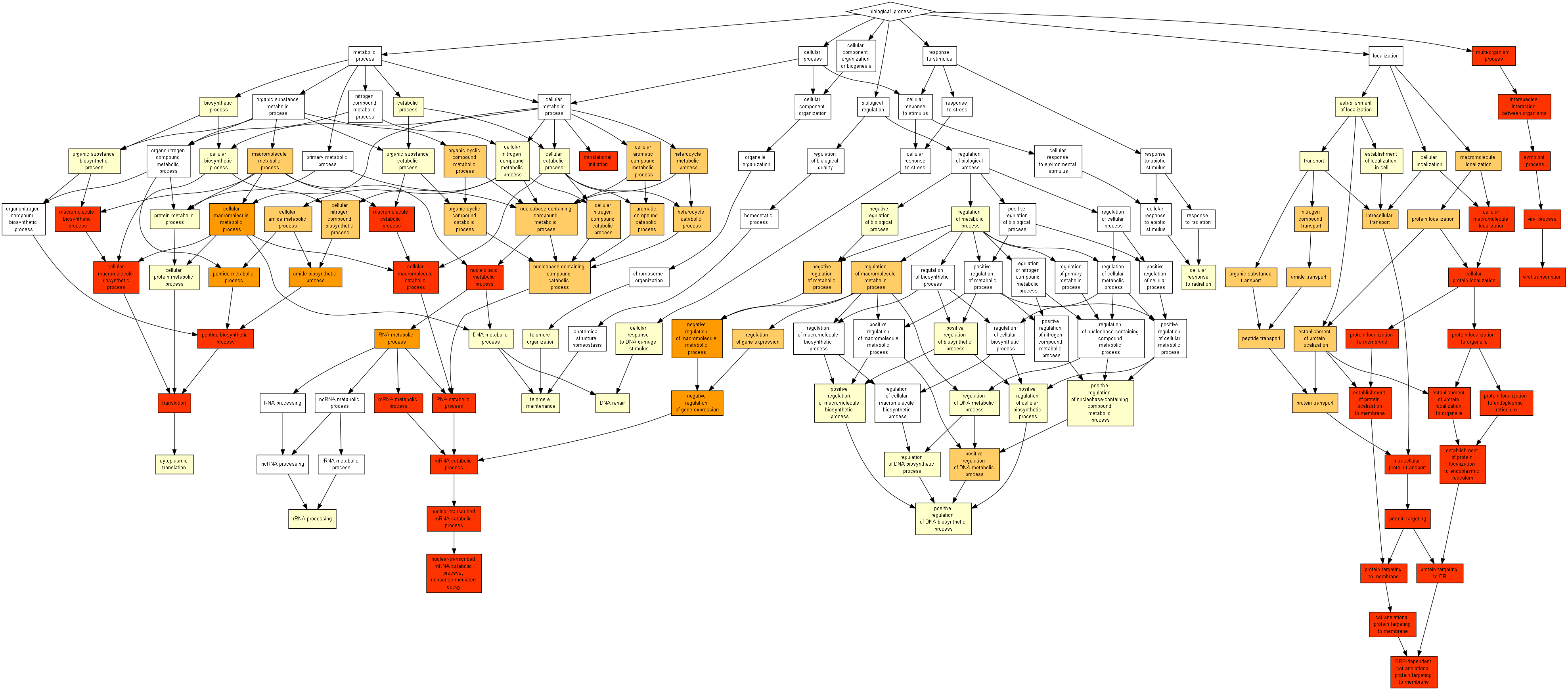
